# RNA-seq mixology: designing realistic control experiments to compare protocols and analysis methods

**DOI:** 10.1101/063008

**Authors:** Aliaksei Z. Holik, Charity W. Law, Ruijie Liu, Zeya Wang, Wenyi Wang, Jaeil Ahn, Marie-Liesse Asselin-Labat, Gordon K. Smyth, Matthew E. Ritchie

**Author notes:** These authors contributed equally (Aliaksei Z. Holik, Charity W. Law).

## Abstract

Carefully designed control experiments provide a gold standard for benchmarking different genomics research tools. A shortcoming of many gene expression control studies is that replication involves profiling the same reference RNA sample multiple times. This leads to low, pure technical noise that is atypical of regular studies. To achieve a more realistic noise structure, we generated a RNA-sequencing mixture experiment using two cell lines of the same cancer type. Variability was added by extracting RNA from independent cell cultures and degrading particular samples. The systematic gene expression changes induced by this design allowed benchmarking of different library preparation kits (standard poly-A versus total RNA with Ribozero depletion) and analysis pipelines. Data generated using the total RNA kit had more signal for introns and various RNA classes (ncRNA, snRNA, snoRNA) and less variability after degradation. For differential expression analysis, voom with quality weights marginally outperformed other popular methods, while for differential splicing, DEXSeq was simultaneously the most sensitive and the most inconsistent method. For sample deconvolution analysis, DeMix outperformed IsoPure convincingly. Our RNA-sequencing dataset provides a valuable resource for benchmarking different protocols and data pre-processing workflows. The extra noise mimics routine lab experiments more closely, ensuring any conclusions are widely applicable.

## Introduction

Transcriptome profiling experiments are widely used in functional genomics research and have helped advance our understanding of gene regulation in health and disease. Throughout the evolution of this technology, from probe-based quantification on microarrays through to sequence-based transcript counting using second and third generation sequencing, researchers have conducted specially designed control experiments to benchmark different platforms and analysis methods. An early high profile example focused on the Affymetrix gene expression platform [1] using a spike-in design and a dilution data set [2]. These experiments became the gold standard for benchmarking different pre-processing algorithms [3] during the rapid development of new background correction, normalisation and transformation methods [4] for the Affymetrix technology. The spike-in design allows bias to be assessed for a small number of RNA molecules that have predictable fold-changes (FCs) when samples with different spike concentrations are compared with one another, while for all remaining genes, no change in expression should be observed. The dilution design on the other hand affects the expression level of every gene in the same way, so that when comparisons between pairs of samples are made, predictable FCs will be induced. This allows bias and variance to be assessed using the data from every gene.

Another popular configuration for control experiments is the *mixture* design, where two distinct samples are mixed in known proportions, inducing predictable gene expression changes across the entire series [5, 6, 7]. This approach is exemplified by Holloway *et al.* (2006) [8], who designed and conducted an experiment to compare a range of microarray platforms. In this study, RNA from MCF7 and Jurkat cell lines were profiled as both pure and mixed samples in different proportions (94%:6%, 88%:12%, 76%:24% and 50%:50%). Holloway *et al.* [8] pioneered an approach in which a nonlinear curve is fitted to the expression values for each gene as a function of the mixing proportions, yielding consensus estimates of signal and noise per gene, allowing comparisons between platforms to be easily made.

The Microarray Quality Control Consortium (MAQC) used this design in a large scale inter-lab (11 sites), inter-platform comparison of 7 microarray technologies using commercially available bulk RNA sources (Universal Human Reference RNA from Stratagene and Human Brain Reference RNA from Ambion) profiled as pure and mixed samples in 2 different proportions (75%:25%, 25%:75%) [9]. This project matched genes between platforms using the probe sequences and looked at reproducibility as measured by the coefficient of variation between replicate samples (*within* and *between* labs), rank correlations between microarray and qPCR platforms and consistency of differential expression results (amongst others). The study concluded that all platforms compared are capable of producing reliable gene expression measurements.

With the advent of RNA-sequencing (RNA-seq), the MAQC project was extended by the Sequencing Quality Control (SEQC) consortium [10] which used the same design to compare different technologies (Illumina HiSeq, Life Technologies SOLiD and Roche 454) across labs (10 sites) using different data analysis protocols (aligners, gene annotations and algorithms for detecting differential expression). In this analysis, the built-in truth from the mixture design was used to measure consistency in 2 different ways (correct titration ordering across the 4 samples and ratio recovery) in order to compare study sites and analysis methods. In addition, spike-in controls enabled assessment of how well changes in absolute expression levels could be recovered. This authors concluded that assessing relative changes in gene expression was far more reliable than absolute expression changes.

Previous mixture experiments performed using either microarray or RNA-seq have a number of well-known limitations. The first is that the samples used are all identical, coming from the same source of bulk RNA, meaning that any variation observed is purely technical in nature. In practice, biological noise is a key source of variability in both microarray [11, 12] and RNA-seq experiments [13] that should ideally be simulated in the experimental design. The second related issue is that sample quality is uniform and high. In regular experiments, both biological variation and variation in RNA quality can be expected.

To address these shortcomings, we designed and conducted a mixture experiment that simulated variability beyond the purely technical. This experiment compared two popular library preparation methods (Illumina’s TruSeq poly-A mRNA kit and Illumina’s TruSeq Total Stranded RNA kit with Ribozero depletion) using short reads (100bp) obtained from the Illumina HiSeq platform. In the sections that follow we present details on the experimental design and quality control of this data set and results from the various methods compared using the inbuilt truth available from the mixture design. Popular differential expression analysis methods, differential splicing algorithms and deconvolution algorithms were compared using these data.

## MATERIALS AND METHODS

### Experimental design and sample preparation

The design of the mixture control experiment ensures the FCs for each gene will follow a predictable dose-response, as initially proposed in Holloway *et al.* (2006) [8]. This design was also used in the MAQC [9] and SEQC projects [10]. A pilot RNA-seq experiment involving 5 cell lines (H2228, NCI-H1975, HCC827, H838 and A549, obtained from ATCC) where RNA from each was profiled in duplicate using the same experimental conditions, library preparation method (Illumina’s TruSeq Total Stranded RNA kit with Ribozero depletion) and analysis pipeline described below. Based on this data the two most similar (NCI-H1975 and HCC827) were chosen for the main study.

To obtain samples for the mixture study, cell lines from a range of passages (2-4) were grown on 3 separate occasions in RPMI media (Gibco) supplemented with Glutamax and 10% fetal calf serum to a 70% confluence. Cell lines were treated with 0.01% Dimethyl sulfoxide (Sigma), and after 6 hours, cells were collected, snap-frozen on dry ice and stored at −80°C until required. Total RNA was extracted from between half a million and a million cells using a Total RNA Purification Kit (Norgen Biotek) with on-column DNAse treatment according to the kit instructions. RNA concentration for each pair of replicates to be mixed was equalised using Qubit RNA BR Assay Kit (Life Technologies) so that both samples in a pair had the same concentration (concentration for all pairs was in the range of 100 ng/*μ*l). Replicates of pure NCI-H1975 (100:0) and pure HCC827 (0:100) and intermediate mixtures ranging from 75:25 to 50:50 to 25:75 (Figure 1) were obtained. We refer to these samples labelled as 100:0, 75:25, 50:50, 25:75, and 0:100 in Figure 1 as *100*, *75*, *50*, *25*, and *0*, respectively in the remainder of the paper.

**Figure 1:**
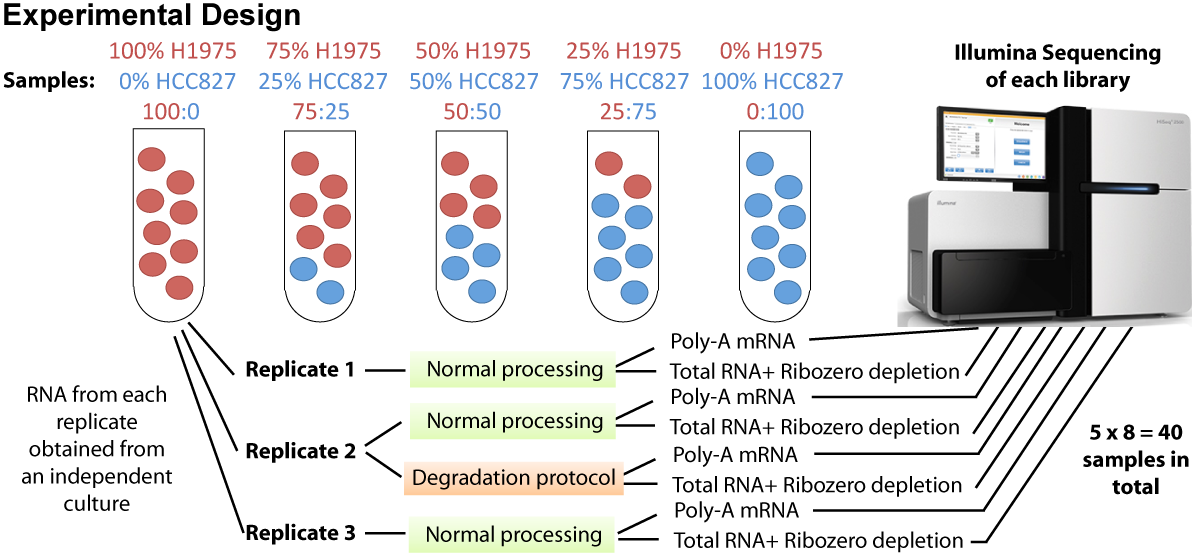
Overview of experimental design of mixture control experiment. RNA from two lung cancer cell lines (NCI-H1975 and HCC827) were obtained after culture on 3 separate occasions to obtain samples for 3 replicates to simulate some degree of biological variability. RNA from each replicate was either kept pure or mixed in 3 different proportions. The second replicate of each mixture was split in two and either processed normally or heat treated (incubated at 37°C for 9 days, see Methods) to degrade the RNA and simulate variations in sample quality. Each sample was then processed using either Illumina’s TruSeq RNA v2 kit (Poly-A mRNA) or Illumina’s TruSeq Total Stranded RNA kit with Ribozero depletion followed by sequencing on an Illumina HiSeq 2500 to obtain 100bp single-end reads for further analysis.

Samples were mixed in these known proportions three times to create three independent replicates of each mixture. All mixtures corresponding to the second replicate were split into two equal aliquots. One aliquot was processed normally (we refer to this as the ‘good’ replicate), while the second aliquot was degraded by incubation at 37°C for 9 days in a thermal cycler with a heated lid (we refer to this as the ‘degraded’ replicate). The RNA Integrity Number (RIN) determined using TapeStation RNA ScreenTape (Agilent) was between 6 and 7 for the degraded samples and around 9 for the good samples (see Supplementary Figure 2). 10 *μ*l (approximately 1 *μ*g) from each replicated mixture (both good and degraded) were used for Next Generation Sequencing library preparation using two different protocols: Illumina’s TruSeq Total Stranded RNA kit with Ribozero depletion and Illumina’s TruSeq RNA v2 kit. Libraries were quantified and normalised by qPCR, as recommended by Illumina, and libraries prepared with the same protocol were pooled together. Library clustering was performed on a cBot with Illumina HiSeq SR Cluster Kit v4 cBot. Each of the two pools of libraries was sequenced as single-end 100 base pair reads over 4 lanes on an Illumina HiSeq 2500 with an Illumina HiSeq SBS Kit, v4. Base calling and quality scoring were performed using Real-Time Analysis (version 1.18.61) and FASTQ file generation and demultiplexing using CASAVA (version 1.8.2). Library quantification, clustering, sequencing, base calling and de-multiplexing were carried out at the Australian Genome Research Facility (Melbourne, Australia).

### Read mapping and counting

FASTQ files from the same libraries were merged and aligned to the *hg19* build of the human reference genome using the Subread and Subjunc (version 1.4.6) software with default settings [14]. Next reads were summarised in various ways according to the NCBI RefSeq annotation (*hg19* genome assembly) using the *featureCounts* procedure [15] in an unstranded manner. The default *featureCounts* behaviour, when generating gene-level counts using the inbuilt annotation, is to count the reads overlapping any of the exons in a given gene. We refer to this annotation as ‘gene-level exon counts’ and use it for the downstream analyses, unless stated otherwise. These counts were used to fit non-linear models of gene expression (see below) presented in Figure 4. In addition to this default annotation, we also generated exon-level counts for the differential splicing analysis (Figure 8). To compare the number of reads mapping to different genomic features between different protocols (Figure 2A), we summarised the reads separately over exons, introns, and intergenic regions based on the inbuilt RefSeq annotation from the Rsubread package. 5’ and 3’ UTR regions were considered as exons and reads were reduced to their 5’ position. To compare the abundance of RNA from different functional categories between protocols (Figure 2B), gene-level exon counts were assigned to different RNA classes based on the Entrez gene type annotation. Gene body coverage plots (Figure 4A) were generated directly from bam files using geneBody coverage.py from the RSeQC [16] software suite based on 3,800 house-keeping genes. To explore gene-level signal in different ways, we ran *featureCounts* using a range of custom annotations including: gene-level exon annotation (default *featureCounts* behaviour - reads overlapping exons are counted and summarised over the entire gene); gene body annotation (reads overlapping any part of the gene body between the transcription start and end sites are counted); conservative gene-level intron annotation (the difference between the full length gene counts and gene-level exon counts, i.e. reads that overlap both an intron and an exon are only assigned to the respective exon). Finally, we also produced a conservative intergenic count. First we produced a combined gene and intergenic annotation, which encompassed the gene body and the intergenic region preceding this gene along the genomic coordinates, regardless of the gene direction. We then counted all reads overlapping each of these amalgamated regions and subtracted the number of reads that overlapped the corresponding gene body, thus obtaining a conservative estimate of the number of reads overlapping the respective intergenic region (i.e. the reads that overlapped both the gene and the preceding intergenic region were only assigned to the respective gene). These counts were used to fit the non-linear models of gene expression (see below) presented in Figures 3-5.

**Figure 2:**
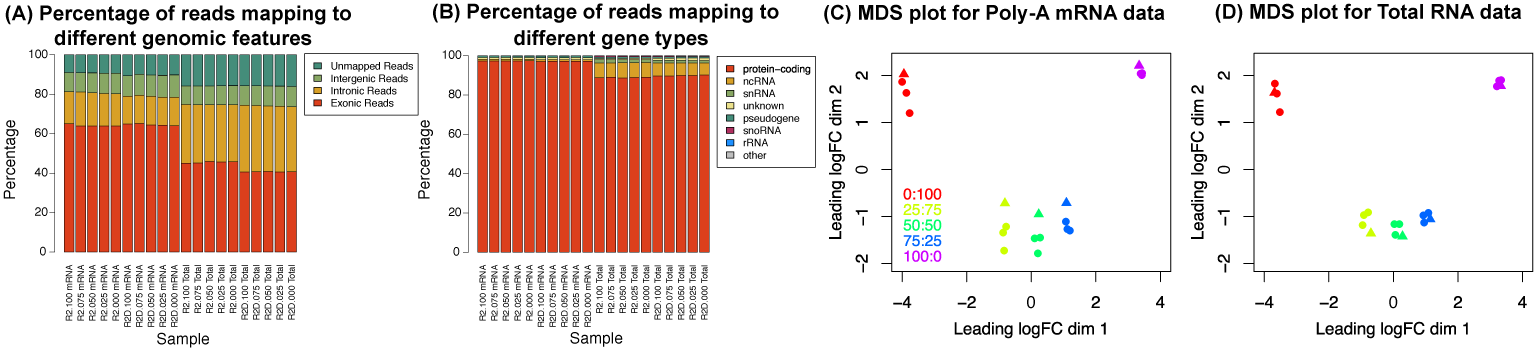
Overview of data quality of mixture control experiment. (A) Mapping statistics of reads assigned to different genomic features for all replicate 2 samples (includes both intact (labels that begin *R2*) and degraded (labels that begin *R2D*) RNA samples). The percentages that could be assigned to exons, introns, intergenic regions or were unmapped are shown in different colours. (B) Mapping statistics of reads assigned to different classes of RNA for all replicate 2 samples. This figure breaks down the gene-level exon reads from panel B according to NCBI’s gene type annotation. Multidimensional scaling plot of poly-A RNA (C) and total RNA (D) experiments showing similarities and dissimilarities between libraries. Distances on the plot correspond to the leading fold-change, which is the average (root-mean-square) log_2_fold-change for the 500 genes most divergent between each pair of samples. Libraries are coloured by mixture proportions, where circles represent good samples and triangles represent degraded samples.

**Figure 3:**
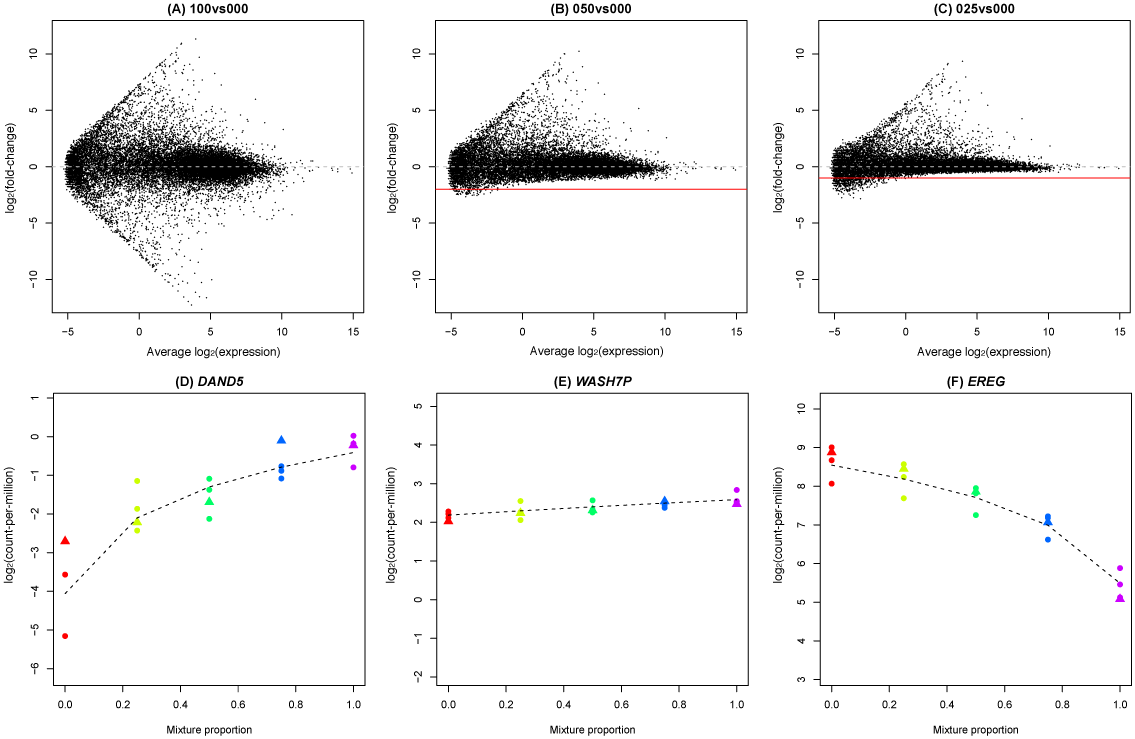
A view of expression changes across the series. Mean-difference plots between the 100vs000 (A), 050vs000 (B) and 025vs000 (C) samples from the total RNA data set show the attenuation of signal as the comparisons get more similar. These mean-difference plots were obtained from the linear model fits from the *voom* analysis of the good samples only across the entire series (15 samples). The solid red line shows the theoretical minimum value that should be possible and the dashed grey line represents log-FC= 0 (no change in expression). Panels D-F show log-CPM values for three genes across the series. The first (*DAND5*, D) has higher concentration in the HCC827 sample, the second (*WASH7P*, E) has approximately equal concentration in each sample and the third (*EREG*, F) has higher concentration in the NCI-H1975 sample. The dashed black line shows the non-linear model fit (Equation 1) obtained using the ‘good’ samples. Libraries are coloured by mixture proportions, where circles represent good samples and triangles represent degraded samples. The concentration parameters and variability estimated from this model can be used to compare analysis methods and protocols.

**Figure 4:**
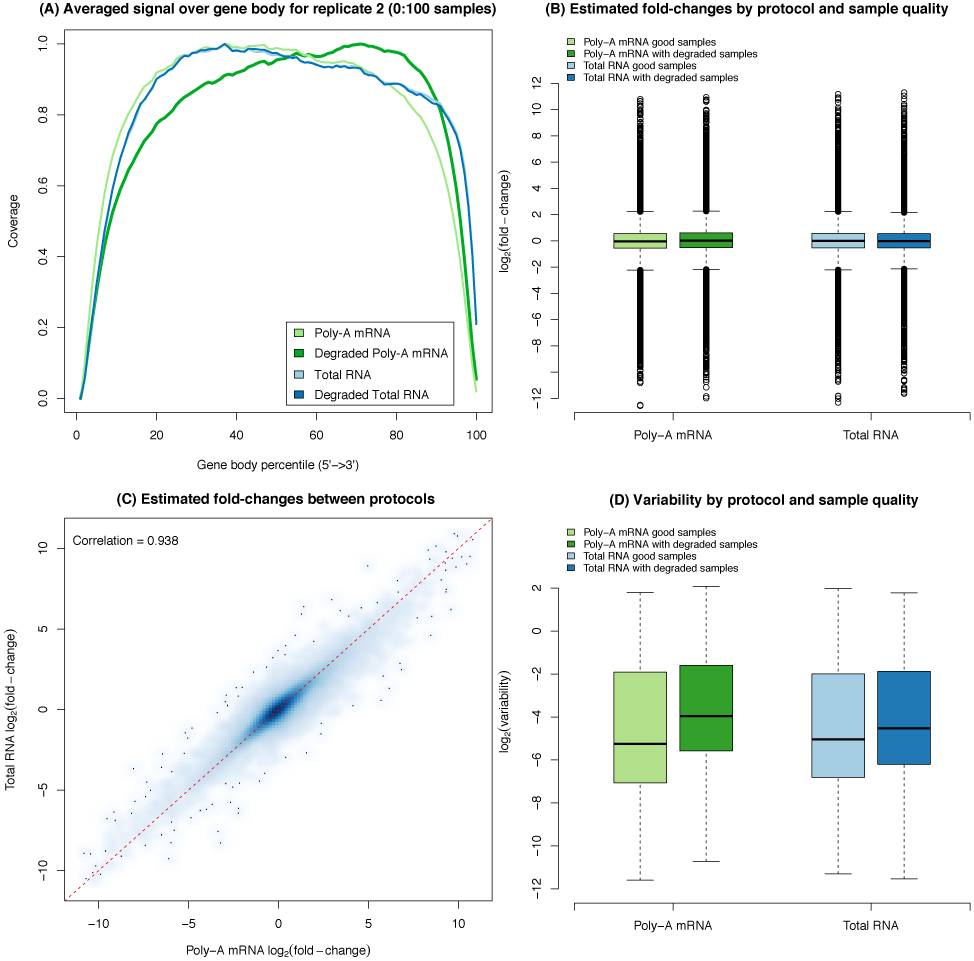
A comparison of library preparation protocols. Plot of read coverage generated by RSeQC for a representative sample from replicate 2 (A). Coverage is fairly uniform for both intact and degraded total RNA samples and for the intact poly-A mRNA sample there is a slightly lower coverage at the 3’ end. For the degraded poly-A mRNA sample, the 5’ end reads are under-represented in this library. (B) Boxplots for the estimated log-FC (*M_g_*) by protocol and degradation state from the non-linear model fit across all genes. Irrespective of protocol and whether degraded samples are included, we see a good dynamic range, indicating that there are no systematic biases between the different protocols. (C) Smoothed scatter plot of the estimated log-FC (*M_g_*) for the total RNA protocol versus the poly-A mRNA protocol based on the good quality samples. The red dashed line represents the equality line. This plot shows good agreement between library preparation methods with no systematic bias. (D) Boxplots of variability (log_2_ 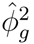) by protocol and degradation state from the non-linear model fit across all genes. As for the read coverage plot, we see that the poly-A mRNA analysis that includes the degraded samples has systematically elevated levels of variability relative to the other analyses.

**Figure 5:**
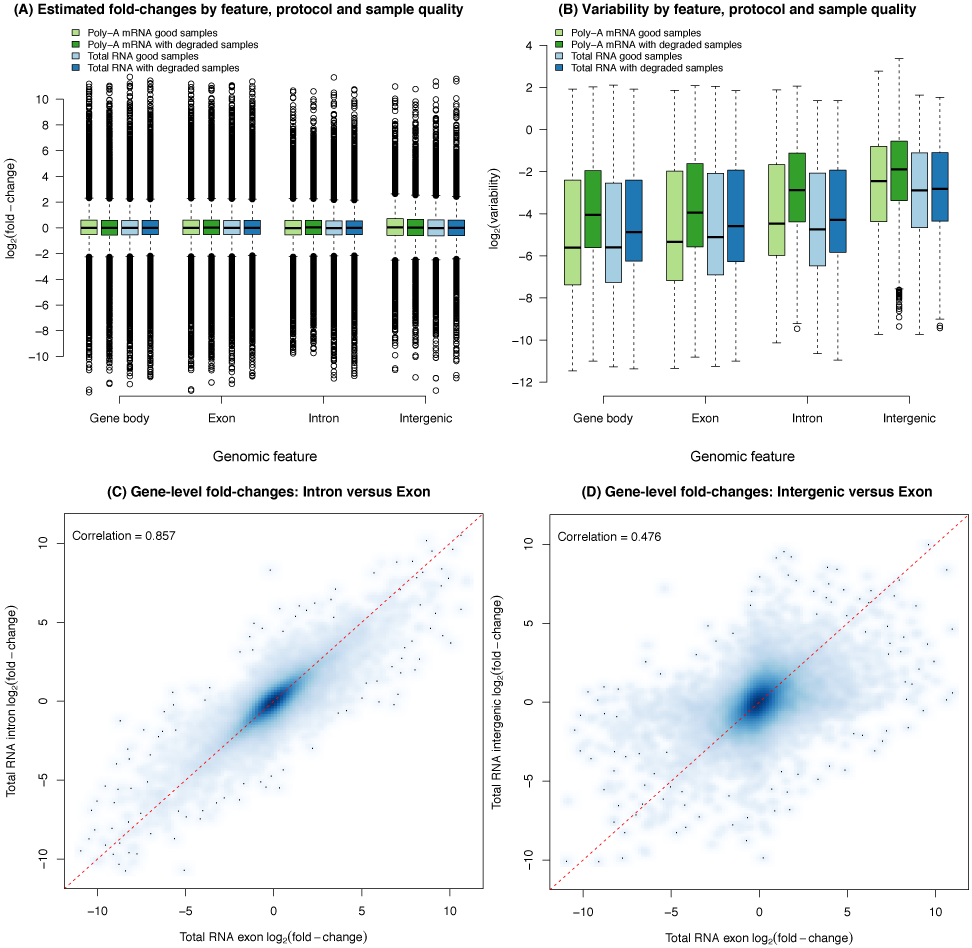
Exploration of signal and noise across different genomic features. (A) Estimated log-ratios (*M_g_*) by protocol and degradation state obtained from nonlinear model fitted to reads that could be assigned to either the entire gene (i.e. overlapping anywhere from the start to the end), or exons only, or the signal left after subtracting the exon-level counts from the gene total, which represents introns or unannotated exons counts, or intergenic regions. Very similar dynamic ranges of log-FC were observed for data obtained from all classes of features. (B) Variability (log_2_ 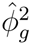) by protocol and degradation state obtained from nonlinear model fitted across the same feature classes shown in panel A. The level of variability from the model fits were generally comparable between counting strategies, although much higher for intergenic counts, which presumably pick up noise. (C) Smoothed scatter plot of estimated log-FC from the non-linear model fitted to the gene-level intron counts (y-axis) versus the log-FCs estimated from the gene-level exon counts for the total RNA data set, which had more intronic reads. (D) Smoothed scatter plot of estimated log-FCs from counts of neighboring intergenic regions (y-axis) versus the log-FCs estimated from the gene-level exon counts for the total RNA data set. The red dashed lines in panels C and D represent the equality line.

### Non-linear modelling of gene expression

We use the nonlinear model described in Holloway *et al.* (2006) to get high precision estimates of the abundance of each gene in the two reference cell lines. For each gene *g*, let *X_g_* be its expression level in NCI-H1975 and let *Y_g_* be its expression level in HCC827. The expression ratio (fold-change) between NCI-H1975 and HCC827 for that gene is therefore *R_g_* = *X_g_/Y_g_*.

For each mixture series, there are 15 RNA samples in total. Write *p_i_* for the proportion of RNA from cell line NCI-H1975 in a particular RNA sample *i*, with 1 − *p_i_* being the proportion of RNA from cell line HCC827, for *i* = 1, …, 15. The expression level of gene *g* in the RNA mix must be *p_i_X_g_* + (1 − *p_i_*)*Y_g_*. The expression FC between the RNA mix and the HCC827 reference must be {*p_i_X_g_* + (1 − *p_i_*)*Y_g_*}/*Y_g_* = *p_i_R_g_* + 1 − *p_i_*, which is an increasing function of *R_g_*. This shows that, regardless of the true values of *X_g_* or *Y_g_*, the expression values for each gene must change in a smooth predictable way across the mixture series.

Write logCPM_*gi*_ for the TMM normalized log_2_-count-per-million value obtained from *voom* for gene *g* and sample *i*. We fit the following nonlinear regression model to the log_2_-count-per-million values for each gene:

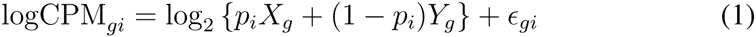

where the *∊*_*gj*_ represent measurement error and are assumed to be independent with mean zero and gene-specific standard deviation *ϕ*_*g*_. The nonlinear regression returns estimates 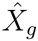, 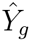 and 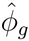 for each gene. The estimates 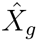 and *Y_g_* are generally more precise than would be obtained from the pure samples alone because they combine information from all the samples. The regression also returns the estimated expression log-ratio, 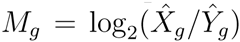, between the two pure samples.

The gene-wise non-linear models were fitted to the matrix of llog_2_-count-per-million values using the `fitmixture` function in the *limma* package. `fitmixture` fits the nonlinear regressions very efficiently to all genes simultaneously using a vectorized nested Gauss-Newton type algorithm [17]. Two separate fits were performed for each gene, the first using only the ‘good’ quality intact samples (15 in total) for each data set, and a second fit to the substituted data set where the good replicate 2 was replaced with the corresponding degraded sample. We repeated this analysis for all possible annotations, i.e. using the log_2_-count-per-million values obtained from gene-level exon, gene-level intron, gene body and intergenic regions.

The nonlinear regression directly estimates the log-FC for each gene between the pure samples. It is also easy to predict what the log-FCs should be between the other RNA mixes. Let the mixing proportion of NCI-H1975 in the first group be *p*, and the mixing proportion of NCI-H1975 in the second group be *q*. Then log-FC for gene *g*, predicted from the nonlinear model, is

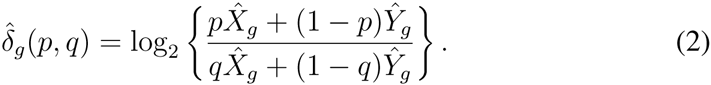

### Differential expression analysis methods

Differential expression analysis is carried out on TMM normalized gene-level exon counts for poly-A mRNA samples. Genes that were expressed in 3 or more samples were kept in the downstream analyses. Genes that fall outside of this criterion are removed. A gene is considered to be expressed if it has a counts-per-million (CPM) value of greater than 1. The number of genes is reduced to 14,981 after filtering on expression.

The counts are normalised using each method’s default, or standard normalisation as described in the respective user guides. Quantile normalisation [18] is carried out for *baySeq* methods; normalisation using the median ratio method [19] is carried out for *DESeq2*; trimmed mean of *M*-values (TMM) method of normalisation [20] is carried out for *edgeR* and *voom* methods.

The *voom* and *voom-qw* methods use default settings in the `voom` and `voomWithQualityWeights` functions, followed by linear modeling and empirical Bayes moderation with a constant prior variance.

Generalised linear models were fitted for *edgeR-glm*, where empirical Bayes estimates of gene-wise dispersions were calculated with expression levels specified by a log-linear model. This differs from *edgeR-classic* where the empirical Bayes method used to estimate gene-wise dispersions is based on weighted conditional maximum likelihood; and where exact tests are carried out for each gene.

For both *baySeq* methods, default settings are used to estimate prior parameters and posterior likelihoods for the underlying distributions. In *baySeq*, counts are modeled under a negative binomial distribution and prior parameters are estimated by the `getPriors.NB` function. In *baySeq-norm*, the underlying distribution of counts are specified as normally distributed; prior parameters are estimated using the `getPriors` function. A default analysis for *DESeq2* is performed using the `DESeq` function.

To obtain mean-difference plots in Figures 3A-C, an analysis of the TMM normalized gene-level exon counts from the total RNA data set using the good samples only (15 in total) was carried out using *voom*, with linear models averaging over the replicate samples and pair-wise contrasts (100vs000, 050vs000 and 025vs000 samples) estimated to get log-FCs (y-axis) and average log-CPM values (x-axis).

To assess the extent of biological versus technical variation in both the good and degraded data sets (using all 15 samples in each) and the pilot SEQC data set from Law *et al.* [21] (16 samples, see separate section below), the *edgeR-glm* pipeline described in Chen *et al.* (2014) [22] was used and the prior degrees of freedom estimated from the empirical Bayes step examined. Plots of the biological coefficient of variation (BCV) versus the average gene expression level were generated using the `plotBCV` function. This linear model analysis was repeated with *voom*, with the prior degrees of freedom from *limma*’s empirical Bayes step reported.

We investigate simple two-group comparisons by subsetting the data for the two groups of interest. For all methods, correction for multiple hypothesis testing was carried out using Benjamini and Hochberg’s FDR approach [23].

For each two-group comparison, the estimated log-FCs were compared with the log-FCs predicted by the nonlinear regression 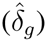. The root-mean-square error in the estimated log-FCs was defined as

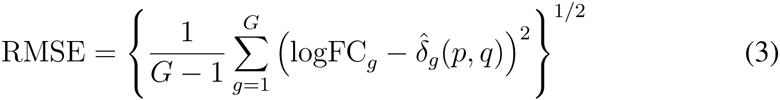

where logFC_*g*_ is the estimated log-FC for gene *g*, *G* is the total number of genes and *p* and *q* are the mixing proportions for the two groups being compared.

### Differential splicing analysis methods

Exon counts were obtained from the poly-A mRNA samples as described in the *‘Read mapping and counting’* section of the Methods. Unexpressed exons are removed from downstream analysis, where an exon is considered to be expressed if it has a CPM value of greater than 0.1 in at least 3 samples. 101,192 exons from 13,974 unique genes remain in downstream analysis.

Counts for the *edgeR* and *limma-voom* analyses are normalised by the TMM method. Whereas counts for *DEXSeq* are normalised using the median ratio method. Like in the DE analyses, simple two-group comparisons are carried out by subsetting the data for the groups of interest and *p*-values are adjusted using Benjamini and Hochberg’s FDR approach [23].

For *DEXSeq*, likelihood ratio tests were performed using size factors and dispersions estimated on default settings. Log-FCs are calculated using the `estimateExonFoldChanges` function and raw gene-wise *p*-values are calculated using the `perGeneQValue` function.

In *voom-ds*, linear modeling on exon-level log-CPM-values were carried out using voom-weights. To test for differential splicing, *F*-tests were performed on each gene using `diffSplice` and `topSplice` functions.

Dispersions (common, trended and gene-wise) in *edgeR-ds* were estimated by calculating an adjusted profile log-likelihood for each gene and then maximising it. In estimating dispersions, the prior degrees of freedom was robustified against outliers. Generalised linear models were fitted with quasi-likelihood tests, where the prior quasi-likelihood dispersion distribution was estimated robustly. Gene-wise tests for differential splicing were carried out using the `diffSpliceDGE` and `topSpliceDGE` functions.

### Deconvolution analysis methods

The two approaches, *DeMix* [24] and *ISOpure* [25], simultaneously estimate the proportion of mixtures and deconvolve the mixed expressions into individual tumor and healthy expression profiles from RNA-seq data. Other published deconvolution approaches thus far do not accomplish these two tasks [26]. Both methods assume a linear mixture structure, that is, *Y* = (1 − *p*)*N* + *pT*, where *Y* is the expression level from a mixed sample, *N* is the expression level from normal tissue, *T* is the expression level from tumor tissue, and *p* is the tumor proportion in the observed mixed sample. *DeMix* is a Bayesian approach that employs the distribution convolution for estimating the proportion of tumor and component specific expressions in tumor-admixed samples.

*ISOpure* uses a Bayesian hierarchical mixture model that further assumes that healthy compartments of tumor-admixed sample can be expressed as the weighted sum of observed healthy samples. Both methods can be fitted using publicly available R [27] functions. Herein the filtering of genes is recommended to exclude genes that a) do not satisfy the linear convolution structure, or b) are uninformative as they have the same expression levels in both sample types such that including them in the model estimation step weakens its ability to differentiate each component.

### Analysis of Pilot SEQC RNA-seq data

The 16 RNA-seq samples (4 samples each from group A, B, C and D) from Law *et al.* (2014) [21] obtained from the pilot SEQC project were downloaded from the voom Supplementary materials webpage (http://bioinf.wehi.edu.au/voom/). This data set was pre-processed by filtering genes with low counts (CPM *>* 1 in at least 4 samples was required), followed by TMM normalization [20]. The `estimateDisp` function in *edgeR* was used to estimate the prior degrees of freedom [22] for this data set. This analysis was repeated using *voom* followed by `lmFit` and `eBayes` in *limma*. The prior degrees of freedom estimated by the empirical Bayes step from each analysis was reported and compared to the mixture experiment described above to assess the relative contribution of biological and technical variation to each data set.

## RESULTS

### RNA-seq mixology balances variability and control

A mixture experiment requires two sources of reference RNA. How to choose these RNA sources has received little attention in the past and, typically, RNA sources that have extremely different expression profiles have been used. Cell lines are good candidates because they provide a replenishable supply of RNA with reproducible characteristics. Our choice of cell lines was guided by a pilot study that looked at gene expression across 5 lung adenocarcinoma cell lines (H2228, NCI-H1975, HCC827, H838 and A549) via RNA-seq. The two cell lines HCC827 and NCI-H1975, which were observed to be most similar according to the multidimensional scaling (MDS) plot (Supplementary Figure 1) and had similar molecular aberrations (both have mutations in *EGFR*: L858R and T790M mutations are found in NCI-H1975 and deletion of exon 19 is observed in HCC827), were selected for the main experiment. The intent here was to have changes that were more subtle than those typically observed in earlier mixture experiments where completely different tissue types, were compared. For example, use of the MCF-7 breast cancer cell line and Jurkat human T lymphocyte cells in Holloway *et al.* (2006) [8], leads to nearly every gene being DE, which is atypical of regular experiments. The experimental design of our study, which consists of 2 pure RNA samples and 3 RNA mixtures each repeated in triplicate, is shown in Figure 1. The cell lines were grown and harvested on three separate occasions to simulate some degree of variation between samples due to lab processing. Each pure RNA sample from a given cell line (denoted as either 100:0 or 0:100) was mixed with a corresponding sample from another cell line in 3 different proportions (75:25, 50:50, 25:75), yielding 3 independent replicates for each pure and mixed sample. The second replicate from each mixture was divided in two, with one of the samples prepared normally (the ‘good’ sample) and the other undergoing heat treatment (incubation at 37°C for 9 days, see Methods) to systematically degrade the RNA (the ‘degraded’ sample). This process was effective, as shown in Supplementary Figure 2, with the RNA integrity number (RIN) between 6 and 7 for the degraded samples, and above 8.5 for the regular samples. Finally, each sample was split into 2 aliquots, which were used to prepare RNA-seq libraries using Illumina’s TruSeq Total Stranded RNA kit with Ribozero depletion (which we refer to as the *total RNA* kit) and Illumina’s TruSeq RNA v2 kit (which we refer to as the *poly-A mRNA* kit). Libraries were sequenced as single-end 100 bp reads on an Illumina HiSeq2500 instrument producing on average 50 million reads per library (range 32 to 94 million). Reads from all samples were mapped to the *hg19* reference genome using the *Subread* alignment software [14]. Mapped reads were assigned using *featureCounts* [15] according to NCBI’s RefSeq *hg19* gene annotation.

### Comparing read distribution by feature type

We used a variety of strategies to annotate the reads depending on the analysis (see Methods for a detailed description of the annotation strategies). To facilitate comparison of the protocols in terms of read distribution across different genomic features, we reduced the reads mapped using the splice aware aligner *Subjunc* to their 5’ position and assigned them to non-overlapping custom annotations restricted to either exon, intron or intergenic regions (UTR regions were considered as exons). Genomic feature mapping statistics for all replicate 2 samples are shown in Figure 2A. Interestingly, the poly-A mRNA and total RNA kits show greatest differences in terms of reads mapping to introns, which come at the expense of exonic reads. The levels are similar between the degraded and good samples within each protocol, with differences of a few percent at most. The poly-A libraries tend to capture mature poly-adenylated RNA which had undergone splicing, while the total RNA protocol captures both mature and pre-messenger RNA alike, which is reflected in the higher proportion of intronic reads in the latter. These results are similar to the percentages reported in Zhao *et al.* (2014) [28], although we observe a slightly higher proportion of intronic reads in our data.

In order to compare the protocols with regard to read distribution across different RNA classes, we used the gene-level exon counts (default *featureCounts* behaviour) obtained from *Subjunc* aligned data and annotated them according to the gene type information available with the annotation. The percentage of reads assigned to each type for all replicate 2 samples is shown in Figure 2B. Consistent with the protocol design, the Total RNA method is able to recover a greater proportion of reads from non-coding (ncRNAs), small nuclear (snRNAs) and small nucleolar (snoRNAs) RNA species compared to the poly-A mRNA kit (see Supplementary Figure 3 for results per RNA class). Across all samples, the percentage of reads mapping to ribosomal RNAs (rRNAs) was marginally lower for the total RNA protocol compared to the poly-A mRNA kit (Supplementary Figure 3), indicating that the Ribozero depletion used to deplete rRNAs in the former sample preparation method was highly effective, as previously reported [29, 28].

To assess data quality experiment-wide, we generated MDS plots from the gene-wise log_2_ counts per million (log-CPM) (Figure 2C-D). This display clearly separates the samples by mixture proportion, with increasing concentration of NCI-H1975 indicated from left to right in dimension 1 and pure samples (100:0 and 0:100) separating from the mixed samples (75:25, 50:50, 25:75) in dimension 2 in both poly-A mRNA and total RNA data sets. For the poly-A mRNA data (Figure 2C), the replicate samples cluster less tightly, with the degraded samples separating slightly from the non-degraded samples for most mixtures. Samples from the total RNA data (Figure 2D) on the other hand tend to cluster more tightly.

### Exploiting signal from the mixture design genome-wide

We next inspect the typical FCs for the total RNA data using a mean-difference plot from 3 pair-wise comparisons (100vs000, 050vs000 and 025vs000) to visualise the attenuation in signal that occurs as the RNA samples compared become more similar. Figure 3A shows the most extreme results, with average log_2_-fold-changes (log-FCs) (y-axis) versus average expression (x-axis) for the pure samples (100:0 versus 0:100, denoted 100vs000). The FCs cover a wide dynamic range and are symmetric about the log-FC = 0 (no change) line. When samples more similar in RNA composition are compared, the log-FCs are compressed and asymmetric (Figures 3B and 3C). Looking at the log-CPM-values at each mixture proportion for 3 representative genes (Figure 3D-F), we see the dose-response across the mixture that varies according to the true RNA abundance of each gene in the pure samples. The gene *DAND5* for example is more abundant in the HCC827 cell line, while *WASH7P* is expressed at a similar level in both samples, and *EREG* is more abundant in the NCI-H1975 cell line. The non-linear model (see Methods, Equation 1, plotted as dashed lines in Figure 3D-F) can be fitted to the log-CPM values from the ‘good’ data (3 replicates for each mixture) or with the degraded sample replacing the non-degraded sample for replicate 2 sample to estimate the true concentration of a particular gene in each mixture, along with the error.

### Total RNA with Ribozero depletion is a better choice for degraded samples

RNA-seq library preparation methods that rely on capturing poly-A RNA species are susceptible to RNA degradation. As RNA gets degraded, the non-poly-adenylated 5’ end of the molecule becomes under-represented. In order to assess the 5’-3’ bias in our libraries, we used the RSeQC [16] software to plot the read coverage over the gene body calculated using 3,800 housekeeping genes in 4 representative libraries. Both total RNA libraries (good and degraded), as well as the good mRNA library followed roughly the same distribution (Figure 4A). As expected, the degraded mRNA library had a drastically different distribution from the others with 5’ end reads being under-represented in this sample. Compared to both total RNA libraries, the good mRNA library also had a slightly lower coverage at the 3’ end. This effect was likely caused by the residual binding of poly-A sequences to oligo-T beads post-fragmentation and subsequent removal of 3’ end fragments when beads were discarded from the reaction.

To assess the bias and precision genome-wide, we used the gene-wise log-ratios (*M_g_*) and log-variance estimates (log_2_ 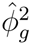) from the non-linear model fit across all genes and generated boxplots of these quantities by library preparation protocol and sample degradation status (Figures 4B and 4D). The signal characteristics are consistent between protocols irrespective of whether degraded samples are included, with a good dynamic range of log-ratios observed (Figure 4B). Moreover, the log-ratios are highly consistent between protocols, with a Pearson correlation of 0.94 (Figure 4C), indicating that no systematic bias is introduced by either kit. Variation on the other hand, was observed to differ between protocols (Figure 4D). The poly-A mRNA analysis that includes the degraded samples was most variable, and the analyses that use only the good quality samples, irrespective of protocol, were the least variable. This is consistent with earlier observations from the MDS plots (Figure 2C-D). For this reason, we chose to focus on the mRNA data in the method comparisons in the remainder of this paper as it shows the most variation due to sample quality and will allow us to compare the performance of methods in the presence of high or low baseline variability by either including or excluding the degraded samples.

### Assessing signal in intronic and intergenic reads

The next issue explored was whether the substantial proportion of reads mapping to introns measured signal or noise. Figure 5 shows the log-ratios (*M_g_*) and log-variances (log_2_ 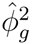) estimated using the log-CPM calculated using 4 different annotations: gene body (i.e. the number of reads overlapping a gene anywhere from the start to the end position, which will include exons (both annotated and unannotated) and introns), gene-level exon (as in Figure 4B), the difference between these two counts in order to obtain a conservative estimate of the intron and unannotated exon counts, or from neighboring intergenic regions that do not contain annotated genes as the control. Very similar dynamic ranges of log-FCs as those observed in Figure 4B were seen from this analysis for each annotation class (Figure 5A). The level of variability from the model fits (Figure 5B) was similar for different gene-centric feature types (gene body, gene-level exon and gene-level intron) with sample degradation increasing the amount of variation in all cases. Interestingly, results from the gene body counting approach are slightly less variable on average than the exon-only results. Results for intergenic regions provide the negative control here, with systematically higher variation than the other feature types. Figure 5C shows a smoothed scatter plot of estimated log-ratios from the non-linear model fitted to the gene-level intron counts (y-axis) versus the log-ratios estimated from the gene-level exon counts (x-axis) for the same gene for the total RNA data set, which had more intronic reads (Figure 2A). The log-ratios showed good agreement, with a Pearson correlation of 0.86. A similar plot displaying the log-ratios obtained from intergenic regions neighbouring a given gene versus the log-ratios from gene-level exon counts has much lower correlation of 0.48 (Figure 5D). While exonic reads are mainly contributed by the more abundant mature mRNAs, intronic signal at the gene-level can presumably be attributed to pre-messenger RNA (pre-mRNA). Since these pre-mRNAs are likely to have a similar relative concentration to their mature counterparts, including these intron reads in the gene-level counts is therefore likely to boost signal, rather than add noise, increasing power.

### Differential expression: 7 methods compared

Differential expression analysis was carried out using 7 popular methods available as part of the Bioconductor project [30]: edgeR [31] using the method of exact tests (*edgeR-classic*) [32] or with generalized linearized models and likelihood ratio tests (*edgeR-glm*) [33]; baySeq [34, 35] with counts modelled either by negative binomial distributions (*baySeq*) or by normal distributions (*baySeq-norm*); limma-voom [36, 21] either with default settings (*voom*) or with sample quality weights (*voom-qw*) [37]; and *DESeq2* [38]. Although all of the above software packages are able to analyze more than two experimental conditions in one analysis, the results we present here are for simple two group comparisons between different mixing proportions. This mimics what is perhaps the most common replicated RNA-seq experimental design in practice, a two group comparison with 3 replicate RNA samples in each group (6 samples in total).

A characteristic of the mixture design is that the same set of genes must be DE between any pair of samples. If a gene is DE between the two pure reference samples, then it should also be DE between any pair of samples with different mixing proportions. However the FCs will be largest when comparing pure samples and correspondingly smaller when comparing intermediate mixtures. Unsupervised clustering of samples confirms that the pure samples have the most extreme transcriptional profiles (Figure 2C-D). It follows that the two group comparison of the pure samples (100vs000) should result in the greatest number of statistically significant DE genes, at any given false discovery rate (FDR). Any two group comparison between other good samples, such as 75 and 25 (075vs025), 50 and 25 (050vs025), or 75 and 50 (075vs050), should ideally result in significant genes that are a subset of those observed in the 100vs000 comparison.

The mixture experiment has no true positives or true negatives for differential expression that are known *a priori*. However we can compare methods by way of their sensitivity, recovery and inconsistency rates. ‘Sensitivity’ can be measured by the total number of DE genes detected. Supplementary Figure 4 shows the number of discoveries made (at a FDR < 0.05 cutoff), which gradually decreases for all 7 methods as the RNA samples compared become more similar in concentration. Using the 075vs025 comparison as an example, ‘recovery’ refers to the proportion of DE genes detected in 100vs000 that are also detected as DE in 075vs025 by the same method, while ‘inconsistency’ refers to the proportion of DE genes in 075vs025 that are not detected in 100vs000 by the same method. Analogous recovery and inconsistency rates can be computed for the 050vs025 and 075vs050 comparisons. These measures examine the performance of each method in the presence of small transcriptional differences, and more specifically, assess how much the power of each method is affected by systematically reducing expression differences. At the same time, we consider the rate of inconsistency to be a reflection of the ‘error’ of a method.

Using a 5% FDR cutoff, *voom-qw* is not only the most sensitive method, but it also achieves the highest recovery rate across all comparisons between good samples (Figure 6A and Supplementary Table 2). For moderate differences in proportions (075vs025), *voom-qw* achieves 82% recovery of DE genes from 100vs000 and 37-39% recovery for subtle differences (050vs025 and 075vs050). Thus *voom-qw* retains much of its power even when transcriptional differences are reduced. Most other methods have recovery rates only moderately lower than that of *voom-qw*, except for *baySeq* and *baySeq-norm* which have much lower recovery rates than the other methods: 62–65% for 075vs050, and 10–14% for 050vs025 and 075vs050. *baySeq* loses power more quickly than the other methods when the transcriptional differences are small.

**Figure 6:**
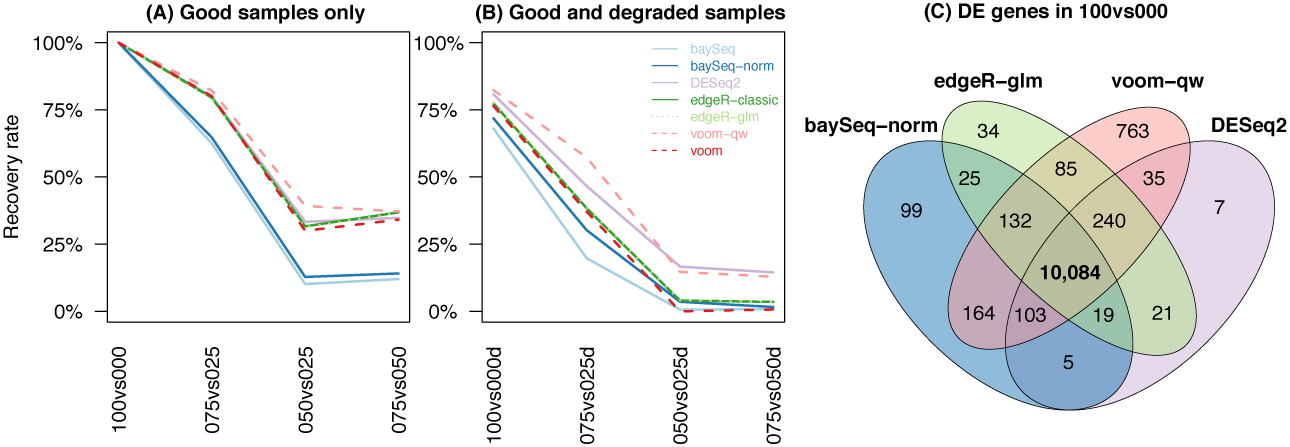
Recovery rate of differential expression methods. The rate at which DE genes are recovered from 100vs000 are displayed for comparisons using good samples only (A) and for those using good and degraded samples (B). Each method is shown in a distinct colour, with different line-types used when results are overlapping to allow them to be distinguished from one and other. The number of common DE genes for 100vs000 across select methods are displayed in (C). For a given software package, only the method with higher recovery is shown.

The experimental design allows us to perform an analysis on the regular samples and compare our results to the analysis that includes the degraded samples. Replacing the non-degraded replicate 2 with the degraded sample results in extra within-group variability that can be observed in the vertical spread of points in the MDS plot in Figure 2C for the poly-A mRNA data. Comparisons with degraded samples will be referred to as 100vs000d, 075vs025d, 050vs025d and 075vs050d respectively. These degraded samples are intended to represent low quality samples that are frequently observed in practice [39].

As expected, increasing within-group variability results in lower recovery rates for all methods. In particular, the recovery rates are 68–82% for 100vs000d as compared to 100vs000 (Figure 6B). The *voom-qw* method has the highest recovery rate for 100vs000d and 075vs025d while *DESeq2* has highest recovery by a narrow margin for the most subtle comparisons 050vs025d and 075vs050d. The two *baySeq* methods again have the lowest recovery rates, with the negative binomial modelling of *baySeq* the lowest out of the 7 methods compared. All methods detect differential expression consistently, with low inconsistency rates observed across all comparisons with or without degraded samples (Supplementary Table 2). For large to moderate proportion differences (100vs000d, 075vs025 and 075vs025d) inconsistency rates sit at between 0-1.7%. For subtle proportion differences (050vs025, 050vs025d, 075vs050 and 075vs050d) inconsistency is between 0-0.3%. Although recovery and inconsistency is assessed for each method relative to itself, it is worth noting that most genes detected as DE in the 100vs000 comparison are common to all methods, with an overlap of 10,084 genes (Figure 6C). The *voom-qw* method detects the most unique DE genes (763) and *DESeq2* detects the least unique DE genes (7).

In addition to within-method comparisons, bias can be examined by looking at differences between the log-FC estimated by each method and those predicted by the non-linear model (*M_g_*) over the entire data set and adjusted according to the relevant sample concentrations for the two-group comparisons (see Methods, Equation 2). Predicted log-FCs are calculated using information from all good samples across the experiment – a total of 15 samples. On the other hand, the logFCs estimated by each differential expression method are based only on the 6 samples included in that particular analysis. For this reason, we assume that predicted log-FCs are more accurate than the estimated log-FCs. *baySeq* and *baySeq-norm* do not estimate gene-wise log-FCs, so were not included in this comparison.

Estimates from *voom* match closest to predicted log-FCs for 100vs000 and 100vs000d, with a strikingly near-perfect match when only good samples are used (Figure 7). The tail-end of log-FCs suggests that *DESeq2* is slightly conservative, where the magnitude of estimated log-FCs tends to be slightly smaller than the predicted values; whilst *edgeR* is slightly liberal, with estimated values slightly larger than predicted values. *edgeR-glm* and *edgeR-classic* produce almost identical log-FC estimates. The concordance between estimated and predicted log-FCs drops in 075vs050 and 075vs050d, when the differences are most subtle. Here, *edgeR-glm* estimates log-FCs marginally better than *voom*, matching closest to the predicted values. Across all comparisons, *DESeq2* estimates are most biased, with the greatest deviation from the predicted values.

**Figure 7:**
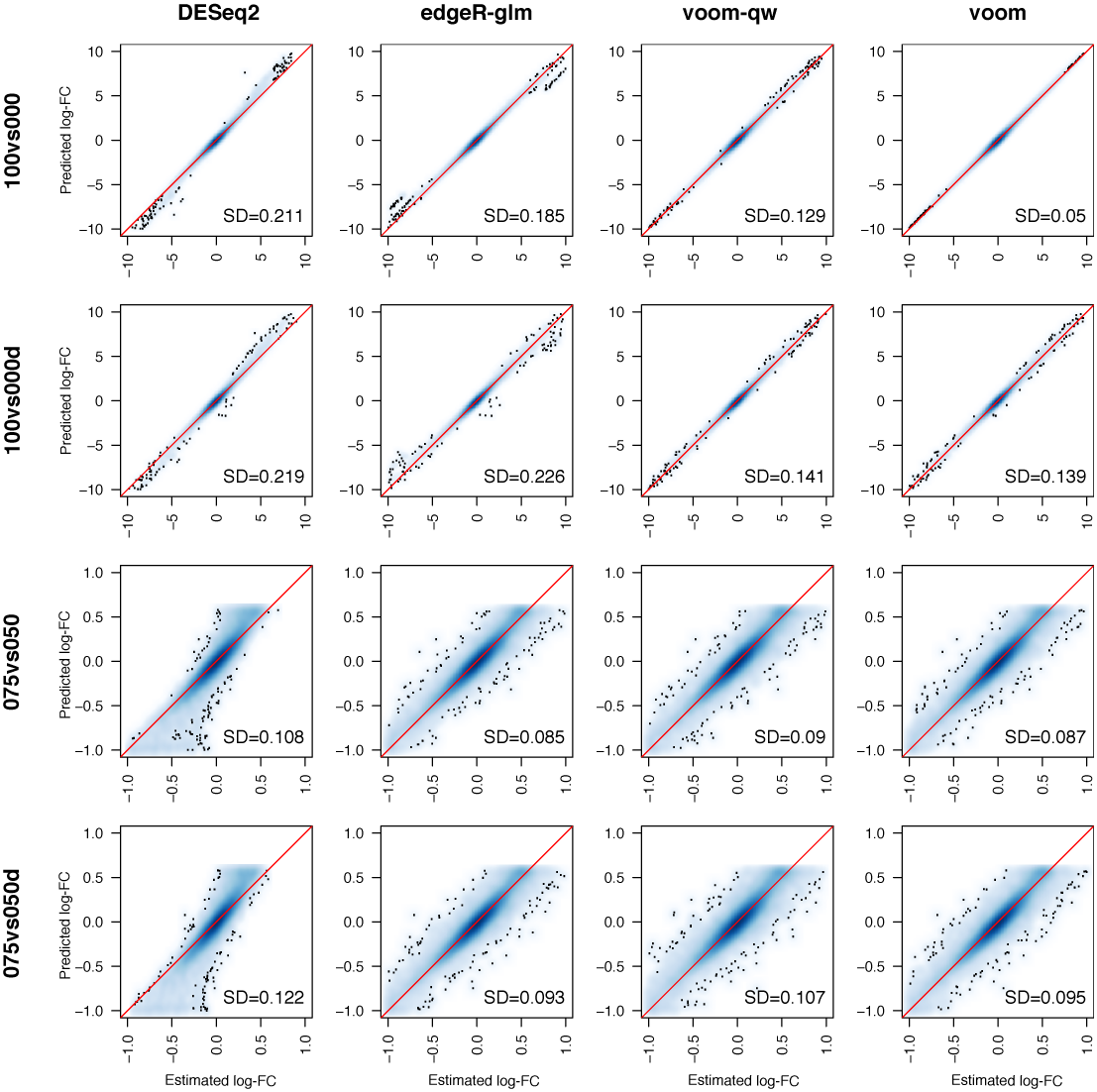
Accuracy of log-FCs estimated by differential expression methods. Gene-wise log-FCs estimated by methods are plotted against predicted log-FCs. Areas with high density of points are shaded in blue, where colour intensity reflects the density of points. Red lines mark equality between estimated and predicted log-FCs. Standard deviations (SD) between estimated and predicted log-FCs are displayed in each panel.

### Comparing variability levels between our mixture and SEQC

Another issue to consider is whether these data sets contain more variability than comparable control data sets that are dominated by technical variation. To assess this, we looked at the prior degrees of freedom obtained from the *edgeR-glm* analysis fitted to all good samples only or the good and degraded samples (15 samples in each analysis). The pilot SEQC RNA-seq data set from Law *et al.* (2014) [21] which consisted of 16 samples (4 samples for A, B, C, and D) was also analysed using *edgeR-glm*. The SEQC data set was pre-processed by filtering genes with low counts (CPM *>* 1 in at least 4 samples was required), then TMM normalized. The prior degrees of freedom was estimated to be 5.30 using the good samples, 4.53 for the good and degraded samples and 67.7 for the SEQC data set. This value measures the consistency of the gene-wise dispersions, with a smaller prior degrees of freedom indicating the values are more gene-specific (i.e. there is more biological variation) and larger values indicating that they are more consistent (i.e. there is more technical variation) and will be more heavily shrunk to a common value. Repeating this analysis with limma *voom* gave similar results with prior degrees of freedom values estimated as 5.29, 4.54 and 45.3 respectively for the 3 data sets. Plotting the biological coefficient of variation (BCV) from the *edgeR-glm* analysis offers another way to visualise this (Supplementary Figure 5). Much greater spread in gene-level variability is observed in the current mixture experiment (Supplementary Figure 5A and 5B) compared to the SEQC data set (Supplementary Figure 5C) in which all genes at a particular expression level have a very similar BCV. This analysis shows that our efforts to simulate additional noise over and above pure technical noise were successful, with our data sets having systematically higher variability than the pilot SEQC data set.

### Differential splicing: 3 methods compared

Detection of differentially spliced (DS) genes was carried out using 3 available Bioconductor methods for the analysis of exon counts: *DEXSeq* [40], *edgeR-ds* [33], and *voom-ds* [36] (see Methods). The recovery rate and inconsistency rate of each method is examined as it was for differential expression. Rate of recovery is defined as the proportion of DS genes detected in 100vs000 that are also detected in another comparison (075vs025, 050vs025 or 075vs050) by the same method. The rate of inconsistency is defined as the proportion of genes detected as DS in a particular comparison (075vs025, 050vs025 or 075vs050) that were not detected in the 100vs000 comparison by the same method.

Using only the good samples, the recovery rates of the 3 methods were similar for each comparison, with *DEXSeq* having slightly higher recovery than *edgeR-ds* or *voom-ds* (Figure 8A). Compared to the differential expression analysis, the differential splicing methods rapidly lose the ability to recover DS genes from the 100vs000 comparison when the samples become more similar – 33-41% recovery for 075vs025, and 2-7% recovery for 050vs025 and 075vs050.

**Figure 8:**
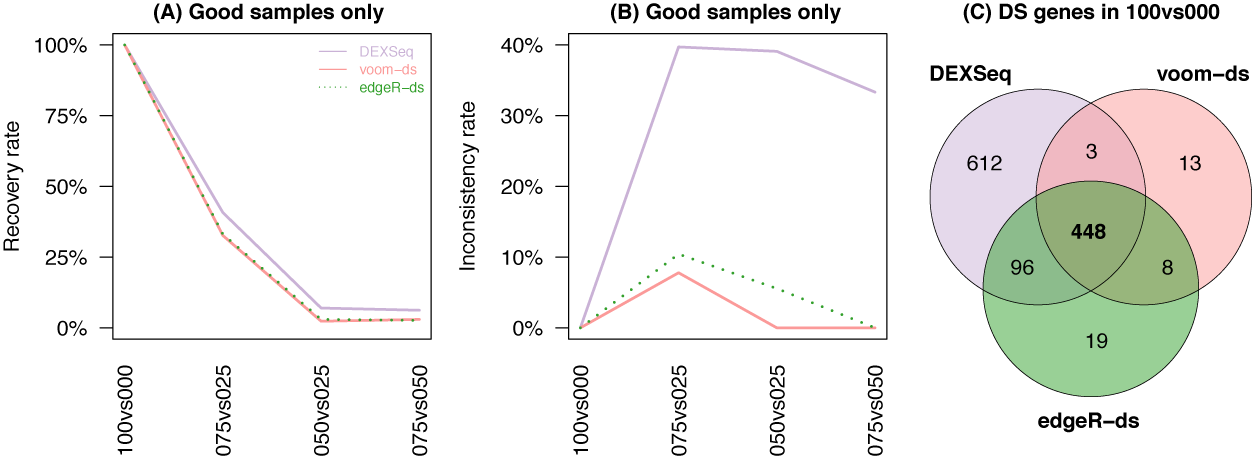
Recovery and inconsistency rate of differential splicing methods. The rate at which DS genes are recovered from 100vs000 (A) and the inconsistency of detected genes (B) are displayed for comparisons using good samples only. Each method is shown in a distinct colour, line-type combination. The number of common DS genes for 100vs000 across the three methods are displayed in panel
C.

Although *DEXSeq* achieves slightly better recovery than *edgeR-ds* and *voom-ds*, this advantage is negated by its very high inconsistency rate (Figure 8B). For the comparisons between subtle proportion differences, methods are expected to have both low recovery and inconsistency rates since methods should be conservative in theory. For comparisons of mixed samples, 33-40% of the genes detected as DS by *DEXSeq* are not detected in its corresponding 100vs000 analysis. This means that a third or more of these DS genes did not achieve significance in the comparison between the pure samples where DS genes can be detected confidently, but were significant when the between sample differences were more subtle.

On the other hand, *voom-ds* and *edgeR-ds* achieve much lower rates of inconsistency, with *voom-ds* being the least inconsistent. For 075vs025, *voom-ds* and *edgeR-ds* have inconsistency rates of 8% and 10% respectively. These rates fall to 0% in *voom-ds*, and 0-6% in *edgeR-ds*, for the 050vs025 and 075vs050 comparisons which are the most subtle. Both of these methods have comparable, but slightly lower recovery rates than *DEXSeq*, however, any gene that is detected as DS by these two methods are likely to be also detected in the 100vs000 comparison. This may suggest that *voom-ds* and *edgeR-ds* have low FDRs.

Overall, *DEXSeq* appears to be liberal relative to *voom-ds* and *edgeR-ds*. For 100vs000, 448 genes are commonly detected by all three methods (37% of the total DS genes identified by any method, Figure 8C), which is much lower than was observed for the differential expression analysis (85%, Figure 6C). Another 612 genes are detected only by *DEXSeq* whereas *voom-ds* and *edgeR-ds* uniquely detect 13 and 19 DS genes, respectively (Figure 8C).

Results from comparisons between the mixture samples (075vs025, 050vs025 and 075vs050) indicate that the differential splicing methods deteriorate more rapidly than the differential expression analysis methods when the expression changes become more subtle, with lower recovery rates and higher inconsistency rates. In general, differential splicing analysis is more complex and challenging than differential expression analysis. Detection of differential exon usage is equivalent to a statistical test of interaction rather than of a simple change in level between two treatment conditions, so there is less statistical power to detect differential splicing. Furthermore, the total gene-level read counts used for differential expression analysis are spread over several exons per gene, leaving much lower counts per feature for exon-level analysis.

### Deconvolution analysis: DeMix versus ISOpure

The mixture design is also ideal for comparing methods that aim to determine the proportions of different sample types present in heterogeneous samples. To this end, we estimate mixture proportions for 12 poly-A mRNA samples using the statistical methods *DeMix* [24] and *ISOpure* [25]. These methods determine the proportions of two different cell populations (typically normal and cancer cells in a patient sample) present in a mixed sample and deconvolve the expression profiles. The input matrix for both methods are read counts from the four pure samples of each type (100, 0) and the twelve mixed samples (4 each of 75, 50 and 25). After filtering based on FC cutoffs of *>* 1.5 or *<* 0.667, only 2,055 informative genes remained. As shown in Table 1, *DeMix* yielded proportion estimates that were closer to the ground truth (differences of less than 10%) while *ISOpure* largely overestimated mixture proportions across all 12 samples. We also compared the deconvolved expression values from the two methods with the four pure samples using Pearson correlation coefficients. The correlations from *DeMix* are all above 0.95 and were generally higher than those obtained from *ISOpure* (Table 1). This analysis clearly demonstrates that *DeMix* is a more reliable method for deconvolution analysis.

**Table 1:** (Top) Mixture proportions estimated from each sample from the plotA mRNA data by the reference free methods *DeMix* and *ISOpure*. The results for the degraded sample is listed last in each cell in *italics* and the values closest to the true values are <u>underlined</u>. In each case, *DeMix* recovers values closest to the true values shown in the bottom row. Pearson correlation coefficients calculated between each of the deconvolved expression profiles mixed samples and the associated pure tumor sample are also shown. The highest correlation coefficients for each sample is again <u>underlined</u>. Of the two methods compared, *DeMix* tends to have higher correlations (closer to 1).

| Method | 25:75 | 50:50 | 75:25 |
| --- | --- | --- | --- |
| *DeMix* (estimated proportion) | 0.311,0.294,0.284,<u>0.247</u> | 0.563,0.544,0.518,<u>0.516</u> | 0.759,0.730,0.751,<u>0.733</u> |
| *DeMix* (correlation) | 0.994,0.986,0.989,<u>0.996</u> | 0.987,0.986,0.959,<u>0.956</u> | 0.964,0.998,0.998,<u>0.964</u> |
| *ISOpure* (estimated proportion) | 0.654,0.673,0.655,<u>0.615</u> | 0.917,0.913,0.908,<u>0.883</u> | 1.00,1.00,1.00,<u>1.00</u> |
| *ISOpure* (correlation) | 0.975,0.971,0.976,<u>0.965</u> | 0.973,0.968,0.946,<u>0.955</u> | 0.959, 0.986, 0.987, <u>0.964</u> |
| True proportion | **0.25** | **0.5** | **0.75** |

## DISCUSSION

For common tasks like differential expression analysis, there have been many comparison studies that have arrived at different conclusions on the ‘best’ method using different data sets (both simulated and experimental) and assessment criteria [41, 42, 21, 38, 43, 44, 45, 46]. Some find *voom* to be best [21, 41], or *edgeR* [42, 45], or *DESeq2* [38, 44] or *edgeR* and *DESeq2* to be equal best [43]. In terms of our main results from comparing DE testing methods on our mixture data set, we find *voom-qw* to be most sensitive followed by *DESeq2*, *edgeR* (glm and classic), *voom* and finally *baySeq* (both normal and regular) is the least sensitive. Recent work has shown that combining differential expression methods in a weighted manner [47] can offer better results than relying on individual methods. Further exploration of this approach on data sets such as this, while including methods such as *voom-qw* would be an interesting topic for future research. Optimising such an approach has the potential to ensure, that for any given data set, the best possible results are obtained.

For differential splicing, our study is the first to compare *DEXSeq* with the competing methods in *edgeR* and *limma*. The latter methods were found to be more conservative than *DEXSeq*, identifying somewhat fewer differential splicing events, but achieved much better consistency than *DEXSeq*, which seems to have a high empirical false positive rate.

Unlike the study of Gallego Romero *et al.* (2014) [48], who degrade PBMC samples by leaving them at room temperature for varying lengths of time, our work couples sample degradation with a classic mixture design, which allows precision and bias to be assessed for every expressed gene. Our experiment has 3 replicate samples per mixture, which is close to the minimum that can be used for tasks like differential expression analysis and differential splicing analysis, so our comparisons should be interpreted with this in mind, in that we are working in the most difficult setting, which is nonetheless a common one in practice. A limitation of the current study is that we simulated a third of the data to be more variable by degrading one sample from each of pure and mixed group. This study design particularly tests the ability of differential expression methods to tolerate heterogeneous variability across the samples. If we turn this around and have the majority of samples being more variable, tasks like detecting differential expression with such a small sample size will become much harder and yield many fewer significant results, irrespective of the method chosen, particularly for the more subtle mixture comparisons.

Further extensions to this experiment could include other library preparation methods, such as exome capture, which has been shown to improve the results from degraded samples [49] or Globin depletion methods [50], although the mixture would need to re-designed to include an appropriate blood sample to make this comparison informative. The inclusion of controls such as the External RNA Controls Consortium (ERCC) spike-in mixes (1 and 2) in different samples is another enhancement that would provide transcripts with known FCs as a further tool for assessing bias. Other genomics applications, such as ChIP-seq or RNA-seq using new technologies, such as long read sequencing platforms from companies such as Pacific Biosciences (Sequel) or Oxford Nanopore Technology (MinION and PromethION) could also benefit from having available a reference data set such as this to allow methodology testing and inter-platform comparisons.

Our analysis has included many of the most popular Bioconductor tools but is not intended as an exhaustive methods comparison. Different algorithms for other common analysis tasks such as read alignment and gene counting, normalization, fusion-detection, intron retention analysis and gene set testing, to name a few possibilities, could also be compared using this data set. We also did not test every capability of the software packages, in particular, we have not tested the ability of the software packages to analyse complex experimental designs with multiple treatment factors and blocking variables. We have also not tested the newer features of some of the packages, for example edgeR’s robust dispersion estimation [51], edgeR’s quasi-likelihood pipeline [52] or limma’s robust empirical Bayes [53]. A natural way to facilitate widespread systematic evaluations of other methods would be to incorporate this data set into one of the recently released RNA-seq benchmarking tools such as *RNASEQcomp* [54] or *RNAonthebench* [55].

## CONCLUSION

We have generated a unique control experiment that is the first to include sample-level heterogeneity induced by systematically degrading samples in order to simulate this routine source of variation. The variability added to our study has shifted the noise profile away from the pure technical end of the spectrum typical of previous mixture experiments much closer to variation levels seen in regular experiments. The classic mixture design allows precision and bias to be quantified via a non-linear model for each gene, and used as a basis for comparing different sample preparation methods. The comparison of poly-A mRNA and total RNA sample preparation kits from Illumina saw differences in the detection of several RNA classes, with more reads mapping to ncRNA, snoRNAs and snRNAs using the total RNA protocol. This comes at a price, with the total RNA protocol sequencing a greater number of intronic reads compared to the poly-A method. Our investigations showed that these intronic reads measure signal rather than random noise irrespective of the library preparation method used. The estimated log-FCs obtained from modelling either intron counts or exon counts at the gene-level were highly correlated, indicating that including these reads in *gene-level* analyses rather than ignoring them as is standard practice offers a simple way to boost signal on the order of 20-30%. This observation warrants further investigation using other published RNA-seq data through re-analysis with and without the intronic reads. It could deliver savings on the cost of sequencing by allowing more samples to be multiplexed per run for an equivalent amount of gene-centric sequencing once exon and intron reads are pooled. The inclusion of degraded samples in the analysis had a more profound effect on the variability of the polyA mRNA data than the total RNA data, which suggests that the latter method is the best choice in experiments where degraded samples are expected, such as in clinical studies.

This mixture design also allows for internal comparisons to be made *within* methods for benchmarking differential expression and differential splicing analysis methods. How well a given method can detect changes when the differences are large and relatively easy to detect is used as the ‘true positive’ set and compared to the results obtained when an independent set of samples which show more subtle changes are compared in a pair-wise manner.

Although most methods perform well at recovering these ‘true positives’ with high overlap in the easiest ‘pure versus pure’ comparison (100vs000), the *voom-qw* method which explicitly deals with sample-level variation slightly outperforms other methods, with *DESeq2* second best, followed by the two *edgeR* methods which perform similarly, then standard *voom*, and finally the two *baySeq* methods which have the least power and consistency.

In terms of bias, *DESeq2* gives FCs that underestimate the consensus values obtained from the non-linear model, indicating that its shrinkage procedure gives conservative results. For differential splicing methods, *edgeR-ds* and *voom-ds* were found to both be conservative relative to the most popular method, *DEXSeq*, however *DEXSeq* tended to have a high empirical false positive rate compared to these other methods based on the within method inconsistency rate.

This is the first time that a mixture experiment has been used to benchmark methods for this application. The results from the deconvolution analysis showed that *DeMix* consistently outperformed *ISOpure*. Our work demonstrates the utility of carefully designed control experiments for benchmarking library preparation kits and analysis methods.

## ACKNOWLEDGEMENTS

We thank Dr Yunshun Chen and Dr Wei Shi for analysis advice and Dr Stephen Wilcox for technical advice on RNA-seq protocols.

## AUTHOR’S CONTRIBUTIONS

AZH performed the experiments. AZH, CWL, RL, ZW, WW, JA and MER performed analysis. AZH, MLAL, GKS and MER designed research. AZH, CWL, WW, JA, GKS and MER wrote the manuscript with the help of the other coauthors.

